# Spatial Attention in the Moving Brain: Dissociable Roles of Neural Alpha Oscillations and Head Rotation

**DOI:** 10.1101/2025.11.10.687554

**Authors:** Malte Wöstmann, Jonas Obleser

## Abstract

The brain is part of a moving body. Most insights about the relationship between neural activity and cognition come from studies involving movement-constrained participants. Here, we test the roles of neural alpha oscillations (~10 Hz) and head movements for auditory spatial attention in human participants (*N*=33). Neural attention filtering showed up as decreased alpha power in the electroencephalogram (EEG) contralateral to targets and vice versa for distractors. Gyroscopic tracking of head movements revealed consistent head turns towards lateral targets. Lateral distractors induced more variable movement patterns. Systematic miniature movements remained even when listeners were instructed not to move. Head rotations improved task accuracy by reducing target uncertainty. Trial-by-trial modulation of neural filtering and head rotation correlated positively, with neural alpha modulation being followed by lateral eye movements and head rotations. Spatial attention emerges from neural coding of relevant and irrelevant stimuli, co-occurring with lateral head rotations supporting sensory sampling.

## Introduction

Attention enables us to process relevant information despite competing distractions ^1–3^. Traditionally, cognitive (neuro)science has focused on uncovering how the mind and the brain implement attention, often treating overt body movements as unwanted noise ^4,5^. However, to understand the basis of cognition, it is necessary to consider that the human brain evolved in a mobile body that can interact with the world ^6,7^. Although it is assumed that attention is not solely the outcome of internal neural computations but realized by overt actions, the mechanistic interplay of neural and motor processes to guide attention is largely unknown.

Movement-constrained laboratory studies have demonstrated the presence of neural attention filters, that is, brain mechanisms associated with target enhancement or distractor suppression ^8–10^. A prominent spatial attention filter is the lateralization of inhibitory alpha oscillations at around 10 Hz, which decrease in brain areas processing targets and vice versa for distractors ^11–14^. But is attention embodied, in the sense that body movements contribute to the success of attentional selection above and beyond what is known on neural mechanisms of attentional filtering? And if so, how does neural filtering relate to overt motor activity?

Several theories have proposed a tight link between attention and motor activity. The premotor theory of attention proposes that spatial attention emerges from the neural systems controlling action, such that attentional shifts rely on the preparation of orienting movements, even in the absence of overt behavior ^15^. Extending this view, the affordance competition hypothesis suggests that potential actions afforded by the environment compete against each other, with attention emerging from the dynamic selection of behaviorally relevant sensorimotor representations ^16^. Building on the concept of active sensing ^17,18^, it is assumed that body movements actively shape sensory processing and contribute to spatial attention by aligning perception with behaviorally relevant locations in the environment.

In the visual modality, strong links between attention and motor activity have been demonstrated. The frontal eye field has been implicated in both, covert shifts of attention and overt orienting via saccadic eye movements ^19^. Furthermore, attentional focusing in memory induces biases in gaze ^20,21^ and head direction ^22^. Unlike many other species, humans cannot actively move their ears, a constraint that has important implications for auditory spatial attention. Yet, electromyographic recordings of the muscles that move the pinna have revealed slight increases in activity on the attended side ^23^. Here, we consider head rotations during auditory spatial attention, which are known to shape auditory scene analysis ^24^ and enhance speech intelligibility in noise ^25^.

Research has demonstrated a connection between neural signatures of attention and motor activity. In animals, strong links exist between body movement and neural dynamics ^26^ and between eye position and auditory neural responses ^27^. Furthermore, motor plans and attention recruit partially overlapping neural resources ^28^. Of relevance to the present investigation, closing the eyes to focus on acoustic input modulates the effect of attention on alpha oscillations ^29^ and oculomotor activity relates to alpha modulation during spatial attention ^30,31^.

Using mobile electroencephalography (EEG) combined with head-mounted gyroscopic tracking, we here examine participants’ head rotation, neural oscillatory activity, and horizontal eye movements during a cued auditory spatial attention task. We hypothesized functional redundancy between lateralized alpha oscillations and head rotation, such that stronger recruitment of one would reduce the need for the other, manifesting as a negative correlation. Contrary, we here show that attention emerges from positive sequential interlocking of alpha modulation with head rotations, with the former reflecting neural coding of ir-/relevant stimuli in space and the latter supporting sensory sampling of target sounds.

## Results

Human participants (*n* = 33) were instructed to attend to one of two spoken numbers, presented simultaneously at two loudspeaker locations (front & left or front & right). The location of the target number was indicated by a fully valid visual spatial cue; the other number served as the distractor (Fig. 1). To equalize overall performance to ~50 % accuracy, the target-to-distractor sound intensity ratio (SNR) was adapted individually before the main experiment (*M*_*SNR*_ = –26.44 dB; *SD*_*SNR*_ = 3.13). No relationship was found between the adapted SNR and the head rotation results reported below. Participants had the task to report the target number on a number pad. The experiment was divided in eight blocks, during half of which participants were permitted to move their heads. We recorded electroencephalography (EEG) and head rotation using a head-mounted gyroscope.

**Figure 1.**
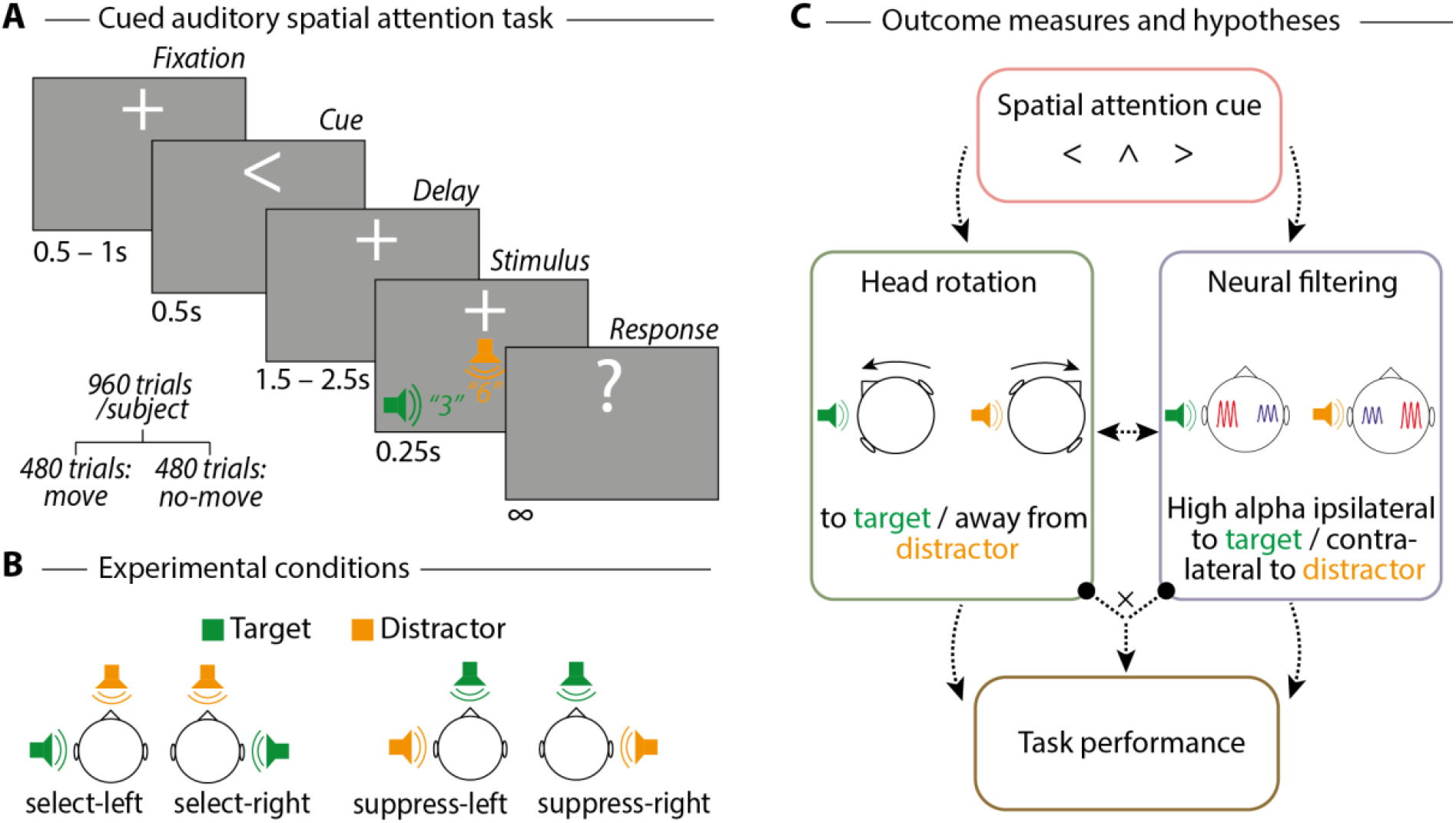
Study design and hypotheses. (**A**) Trial design of the cued auditory spatial attention task. (**B**) Experimental conditions (adapted from^13^). (**C**) Schematic illustration of main outcome measures (head rotation and neural attentional filtering through lateralization of alpha oscillations). Dashed arrows indicate hypothesized relations tested in this study.

### Top-down attention cues induce head rotation

First, we asked whether participants would systematically employ head rotations during spatial attention. When head motion was permitted, cues indicating upcoming targets on the left or right side induced substantial head rotations towards the target (Fig. 2B; cluster *p*-value < .001; *d* = 0.832; cluster time window 0.23–2.08 s). Critically, we found an analogous, if markedly more low-amplitude, pattern of head rotation to lateral targets when movement was not permitted (cluster *p*-value = .001; *d* = 0.65; cluster time window 0.35–1.13 s). These results indicate that attention-related head movements occur involuntarily, even under instructions to suppress overt actions, as is common in conventional, movement-restricted neuroscientific paradigms.

**Figure 2.**
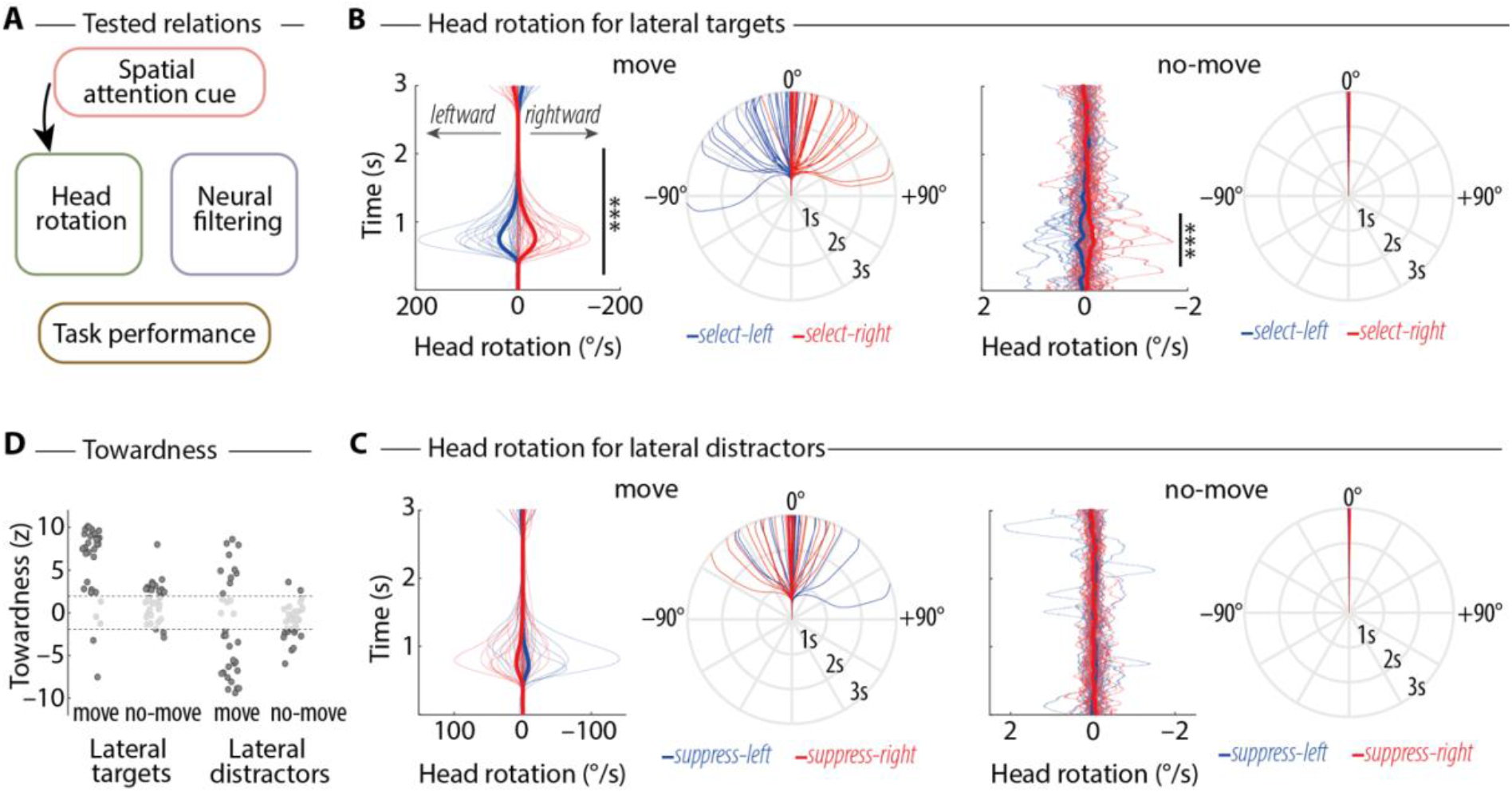
Head rotation during auditory spatial attention. (**A**) Overview of tested hypothesis (black arrow). (**B**) Line plots: Thin and thick lines show respective single-subject and average angular velocity of head rotation (around the yaw axis) for trials with (*move*; left) and without movement permitted (*no-move*; right). Black vertical lines and asterisks indicate significant differences for select-left versus select-right (*p* < .05; cluster-based permutation tests). Polar plots: Illustration of head orientation, calculated by integrating angular velocity over time. Note that absolute values of head rotation/orientation might be slightly underestimated due to tilt of the gyroscope, which was attached to the subjects’ back of the head. (**C**) Same as B but for conditions with lateral distractors. (**D**) Towardness, calculated as the participant-specific *z*-contrast of angular velocity towards versus away from lateral targets/distractors (time window 0.35–1.13 s, i.e., overlap of significant clusters in B). Participants with no significant head rotation (i.e., *–*1.96 < *z* < 1.96) are masked. Note the bimodal distribution in the condition with lateral distractors and movement permitted.

For lateral distractors, even though on average there was a trend for participants to rotate their heads away from the source of distraction, this pattern was not significant on the group level (Fig. 2C; move: all cluster *p*-values > .2; no-move: all cluster *p*-values > .5). However, this absence of a consistent group-level effect was not due to a general lack of head rotations when faced with lateral distractors. Instead, when movement was permitted, 18 out of 33 participants significantly rotated their heads away from the distractor, while 10 out of 33 participants significantly rotated their heads toward the distractor (Fig. 2D). This disparity suggests the use of different strategies to handle lateral distraction (for correlations of head rotation between conditions, see Fig. S1).

### Head rotation improves task performance

Next, we asked whether and to what extent task performance would benefit from the permission to move the head. Across conditions, the proportion of correct responses was higher in blocks with (*M* = 0.578; *SE* = 0.022) versus without movement permitted (Fig. 3B; *M* = 0.545; *SE* = 0.021; main effect movement: *F*_1,32_ = 35.5; *p* < 0.001; *η*_*p*_^*2*^ = 0.526). This amounts to an average movement benefit of 3.27 percentage points^32^. Head motion was not equally beneficial across experimental conditions: While a movement benefit was absent in trials with the attentional target on the right side (*t*_32_ = 0.18; *p* = 0.856; *d* = 0.032; *BF*_10_ = 0.19), it was present for all other spatial arrangements of target and distractor (select-left: *t*_32_ = 3.05; *p* = 0.005; *d* = 0.532; *BF*_10_ = 8.62; suppress-left: *t*_32_ = 3.77; *p* < 0.001; *d* = 0.657; *BF*_10_ = 46.61; suppress-right: *t*_32_ = 3.26; *p* = 0.003; *d* = 0.568; *BF*_10_ = 13.86). Furthermore, the permission to move enhanced confidence and reduced effort ratings (Fig. S2).

**Figure 3.**
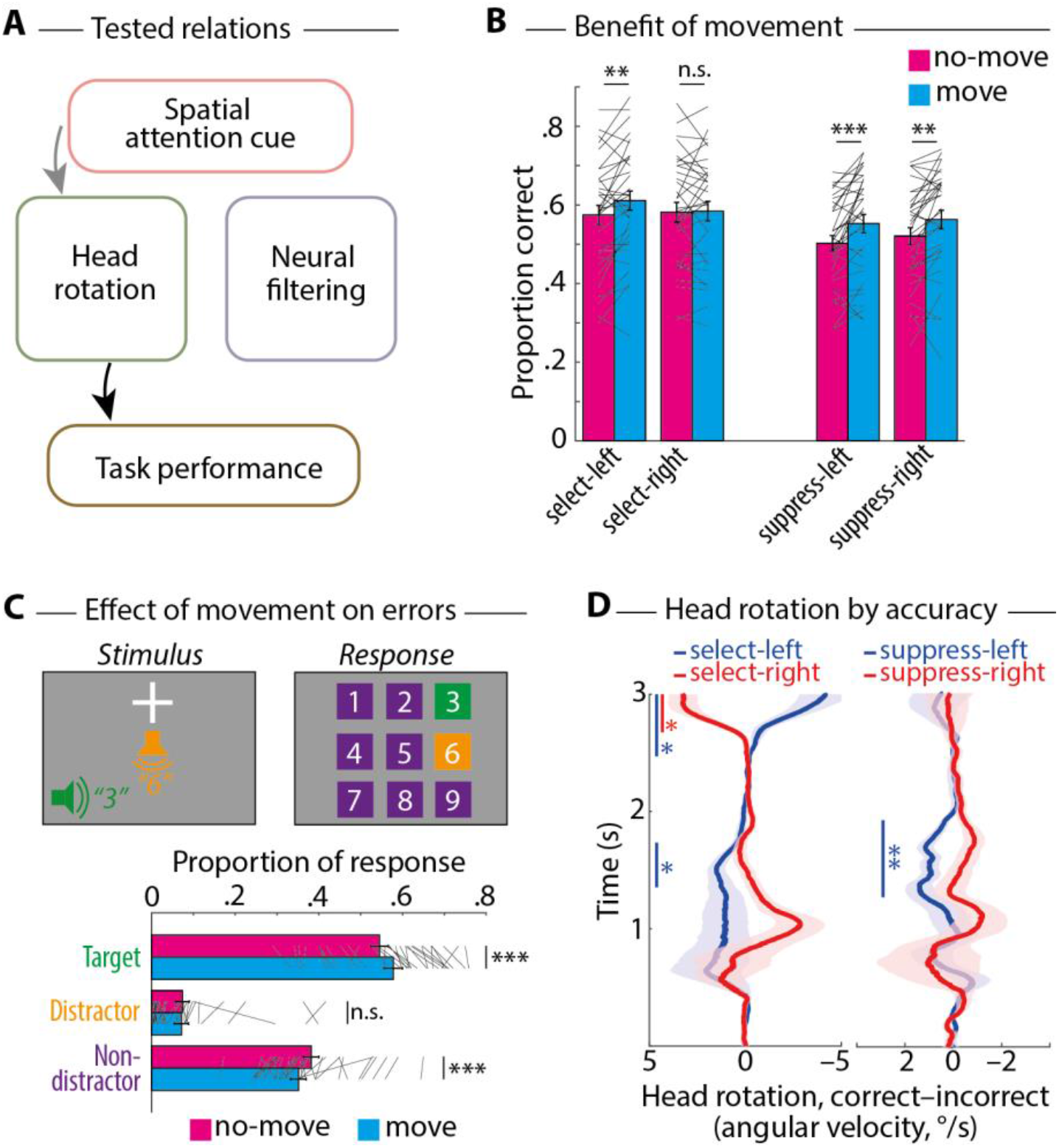
Relation of head rotation and task performance. (**A**) Overview of tested hypothesis (black arrow) and the previously established link (grey arrow). (**B**) Thin lines, bars and error bars show respective single-subject and average of proportion correct ±1 SEM for different conditions. Black lines and asterisks indicate significantly higher accuracy for move versus no-move. (**C**) Grey boxes illustrate three different response types for a given trial (target, distractor, non-distractor). Thin lines, bars and error bars show respective single-subject and average proportion of response types ±1 SEM. (**D**) Lines and shaded areas show respective average angular velocity ±1 SEM for correct minus incorrect trials in conditions with movement permitted (for conditions without movement permitted, see Fig. S4). Vertical lines and asterisks indicate significant differences for correct versus incorrect trials, separately for targets/distractors on the left and right side (*p* < .05; cluster-based permutation tests). * *p* < .05; ** *p* < .01; *** *p* < .001; n.s. not significant.

Independent of head movement, participants performed better in trials with lateral targets than lateral distractors (*F*_1,32_ = 30.74; *p* < 0.001; *η*_*p*_^*2*^ = 0.49), which agrees with previous research^13,33^.

Having established head movement-related benefits for task performance, we next investigated *how* this movement benefit comes about. In our task, the proportion of erroneously reported distractors reflects attention capture by the irrelevant distractor. The proportion of erroneously reported non-distractors reflects target uncertainty (Fig. 3C). Including this factor of error type in the statistical analysis revealed a significant 4-way interaction with movement (no-move vs. move), loudspeaker setup (front & left vs. front & right), and role of lateral speaker (target vs. distractor; *F*_1,32_ = 4.71; *p* = 0.038; *η*_*p*_^*2*^ = 0.128). Post-hoc tests revealed that the permission to move the head did not reduce distractor responses (*t*_32_ = –0.87; *p* = 0.39; *d* = 0.151; *BF*_10_ = 0.26) but non-distractor responses (*t*_32_ = –4.59; *p* < 0.001; *d* = 0.799; *BF*_10_ = 374.3). This movement-induced reduction of target uncertainty was consistent across conditions except for select-right trials, where movement was not beneficial (Fig. S3).

Finally, we asked whether patterns of head rotation would differ for correct versus incorrect trials when movement was permitted. For trials with targets on the left, we found evidence for larger leftward head rotation in correct versus incorrect trials (Fig. 3D; cluster *p*-value = .045; *d* = 0.51 cluster time window 1.35–1.73 s). In addition, head rotation back to the center at the end of a trial was stronger for correct versus incorrect trials with targets on the left (cluster *p*-value = .014; *d* = 0.603 cluster time window 2.47–3 s) and on the right (cluster *p*-value = .02; *d* = 0.748; cluster time window 2.68–3 s). For trials with distractors on the left, participants rotated the head more to the left (i.e., in the direction of the distractor) when they reported the target number correctly versus incorrectly (cluster *p*-value = .002; *d* = 0.54; cluster time window 1.27–1.92 s). In sum, these findings provide evidence for the existence of the hypothesized link between head rotation and task performance.

### Neural signatures of attentional filtering also occur during free movement

Next, we tested whether the lateralization of alpha oscillations—a well-established neural signature of spatial attention in traditional, movement-constrained laboratory studies—would prove to be a robust metric of attention in conditions that closely resemble real life, including head movement.

Well in line with the published literature (see Discussion), cues indicating a lateral target on the left or right side induced higher ipsi-than contralateral alpha power. Notably, however, this finding here proved robust against the permit to move the head (Fig. 4B): Cluster-based permutation tests revealed significant alpha lateralization for no-move and move conditions in similar time windows (no-move: cluster *p*-value = .002, *d* = 1.077; cluster time window 0.54–1.22 s; move: cluster *p*-value = .008, *d* = 0.794; cluster time window 0.42–1.22 s).

**Figure 4.**
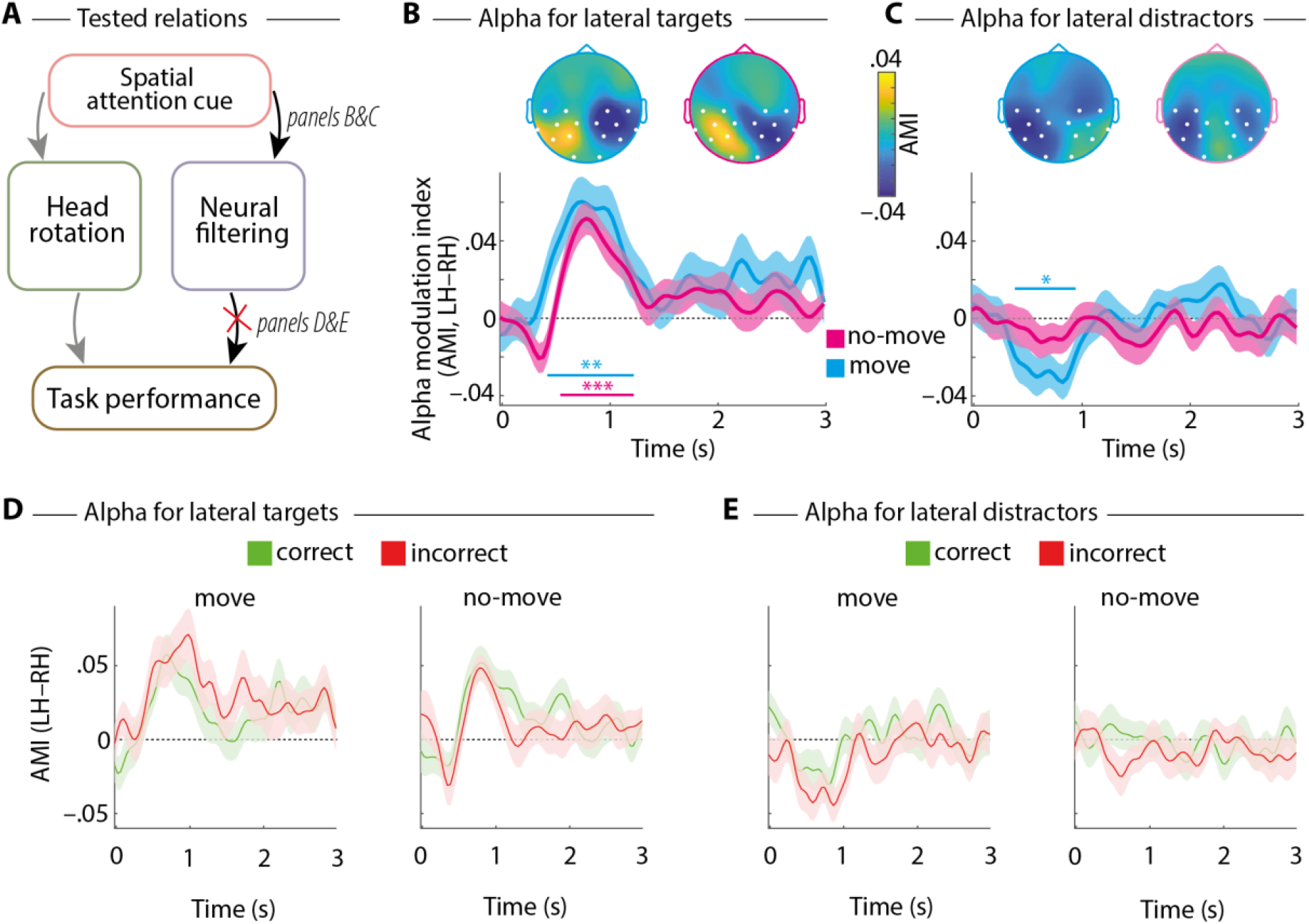
Lateralized alpha oscillations during auditory spatial attention. (**A**) Overview of tested hypothesis (black arrows) and previously established links (grey arrows). (**B**) The Alpha Modulation Index (AMI) for lateral targets was calculated on alpha (8-12 Hz) oscillatory power as: AMI_selection_ = (select-left – select-right) / (select-left + select-right). Lines show the average difference of AMI for the left – right hemisphere (LH–RH), separately for conditions with (blue) and without movement permitted (pink). Shaded areas show ±1 SEM. Horizontal lines and asterisks indicate significant temporal clusters for testing AMI against zero (*p* < .05; cluster-based permutation test). Topographic maps show AMI in time windows of significant clusters. White circles highlight electrodes used for calculating the hemispheric difference of AMI (LH–RH). (**C**) Same as B but for conditions with lateral distractors: AMI_suppression_ = (suppress-left – suppress-right) / (suppress-left + suppress-right). Note that the topographic map for the no-move condition (where no significant cluster was found) refers to the same time interval as the significant cluster for the move condition. * *p* < .05; ** *p* < .01; *** *p* < .001. (**D&E**) Same as B&C but separately for correct (green) and incorrect trials (red). No significant clusters for the difference between correct and incorrect trials were found (all *p* > .5).

For trials with lateral distractors, alpha lateralization showed a significant reversed pattern (Fig. 4C; higher contra-than ipsilateral alpha power) for conditions with movement permitted (cluster *p*-value = .013, *d* = 0.64; cluster time window 0.38–0.94 s) but not for conditions without movement permitted (no clusters found). Importantly, alpha lateralization for lateral targets and distractors did not differ for no-move versus move conditions (all cluster *p*-values > .2), which demonstrates robust neural filtering in moving participants.

To test whether neural filtering would relate to improved task performance, we compared alpha lateralization for correct versus incorrect trials (Fig. 4D & E). No significant clusters were found (all cluster *p*-values > .5), which demonstrates that contrary to head rotation, neural filtering did not relate directly to performance.

### Relation of neural filtering and head rotation

Having established that cued top-down attention cues induce both neural filtering and head rotation, we ultimately asked whether and how the two relate. For trials with lateral targets and movement permitted, alpha power lateralization increased with larger magnitude of head rotation (Fig. 5B, left; movement bin x hemisphere interaction: *F*_1.7,55.5_ = 8.37; *p* = 0.001; *η*_*p*_^*2*^ = 0.207; the interaction was not significant in other conditions, all *p* > .4). The same relationship showed up at the between-subject level, as individuals who on average performed larger head rotations exhibited more pronounced patterns of lateralized alpha oscillations (Fig. 5B, right; Spearman’s *rho* = .413; *p* = .0175).

**Figure 5.**
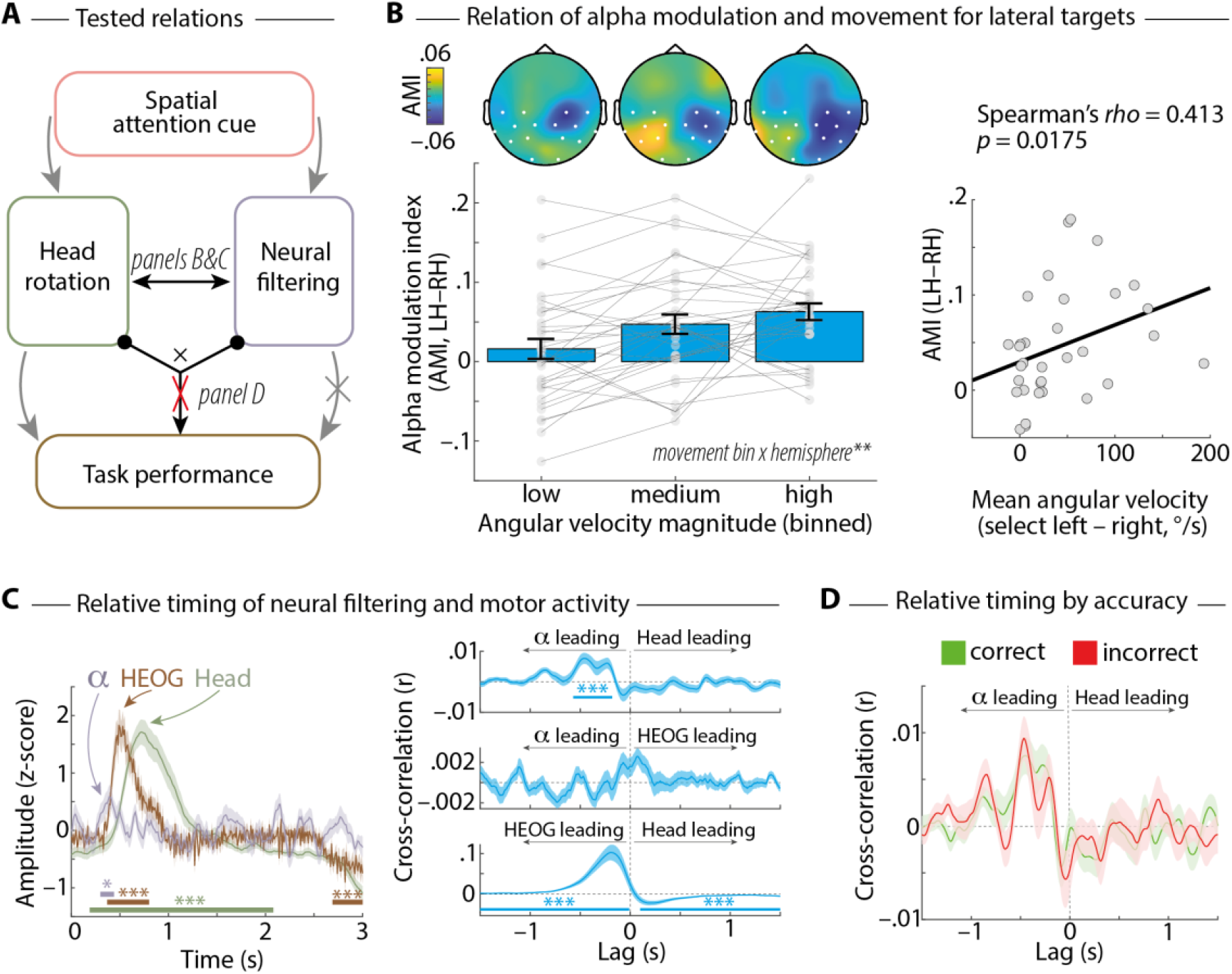
Relation of neural filtering and head rotation. (**A**) Overview of tested hypothesis (black arrows) and previously established links (grey arrows). (**B**) Bars, error bars, and lines show respective mean ±1 SEM and single-subject alpha modulation index (AMI) for the left – right hemisphere (LH–RH), binned for the magnitude of angular velocity (time window AMI: 0.42–1.22 s; time window head rotation: 0.35–1.13 s). Data are shown for trials with lateral targets and movement permitted. Topographic maps show the average AMI, binned for angular velocity. ** Significant movement bin x hemisphere interaction (*p* < .01). The scatterplot shows individual participant’s AMI (LH–RH; time window: 0.42–1.22 s) by mean angular velocity (time window: 0.35–1.13 s), including the least-squares regression line. (**C**) Left: For conditions with lateral targets and movement permitted, we calculated single-trial neural filtering (1^st^ temporal derivative of AMI_single-trial_), head rotation (angular velocity), and the horizontal electrooculogram (HEOG; 1^st^ temporal derivative of amplitude difference at electrode F7–F8). For visualization, since all three measures are supposed to reverse amplitude depending on the side of the lateral target, they were multiplied by –1 in select-right trials, followed by averaging across trials and z-scoring. Lines and shaded areas show mean ±1 SEM, respectively. Horizontal lines and asterisks indicate significant clusters for tests against zero (*p* < .05; cluster-based permutation test). Right: Mean ±1 SEM of single-trial pairwise cross-correlations. Horizontal lines and asterisks indicate significant clusters for tests against zero. (**D**) Same as C (right, upper panel) separated for trials with correct (green) versus incorrect responses (red), which did not differ significantly (all cluster *p*-values > .9). * *p* < .05; ** *p* < .01; *** *p* < .001.

To investigate the temporal sequence of neural filtering and head rotation, we extracted the change in single-trial modulation of lateralized alpha power (i.e., first temporal derivative of left-versus right-hemispheric alpha amplitude) and the change in single-trial head position (i.e., angular velocity). Lateral eye movements have been associated with alpha power modulation ^30,31^ and typically precede head rotation ^34,35^. Despite the absence of eye-tracking data to assess microsaccades in the present study, the change in the single-trial horizontal electrooculogram (HEOG; i.e., first temporal derivative of amplitude difference at electrode F7–F8) was extracted to quantify lateral eye movements (Fig. 5C, left). Descriptively, on the group-level, changes in alpha-lateralization peaked first (~390 ms), followed by changes in HEOG (~490 ms) and changes in head orientation (~720 ms).

As per temporal cross-correlation at the single-trial level, the positive relation of lateralized alpha activity and head rotation was driven by stronger head rotation following stronger neural filtering with a temporal delay of ~375 ms (Fig. 5C, right; cluster *p*-value < .001; *d* = 0.751; cluster time window –0.57 to –0.18 s). This temporal delay was found invariably for correct and incorrect trials (Fig. 5D; all cluster *p*-values > .9 for potential accuracy differences).

Lastly, the data revealed a strong temporal co-occurrence of HEOG and head rotation, with eye-related activity preceding the head-related activity (Fig. 5C, positive cluster: *p*-value < .001; *d* = 0.924; cluster time window –1.5 to –0.01 s; negative cluster: *p*-value < .001; *d* = 0.889; cluster time window 0.1 to 1.5 s). Critically, no significant cross-correlation of lateralized alpha activity and HEOG was found (all cluster *p*-values > .09), which makes it unlikely that the observed relationship between alpha oscillations and head rotation is mediated by lateral eye movement activity.

## Discussion

In this study, we demonstrate that head rotations pose a powerful mechanism co-occurring with auditory attention, above and beyond the well-studied attentional modulations of brain activity. First, head rotation improves task accuracy by reducing uncertainty about the attentional target while neural filtering does not. Second, head rotation is ubiquitous, as listeners perform miniature head rotations to lateral targets even when instructed not to. Third, head rotation relates positively to temporally preceding modulation of lateralized alpha oscillations, suggesting that spatial attention emerges from lateralized alpha oscillations coding the spatial position of ir-/relevant stimuli in space and head rotations to enhance sensory sampling of target sound.

### Auditory spatial attention induces target-directed head rotation

Previous research has shown that listeners’ head movements benefit auditory scene analysis and speech comprehension in noise ^24,25^. Acoustic differences of competing sounds arriving at the two ears facilitate detection and identification – a phenomenon called *binaural unmasking* ^36,37^. Head movements modulate binaural unmasking, with rotations of ~30° being acoustically optimal for lateral targets (90°) and noise in front (0°) ^38,39^. Additionally, movement might support cognitive offloading ^40^, that is, the use of physical action (i.e., head rotation) to reduce cognitive demand (i.e., keeping track of where to focus attention). Here, we demonstrate that preparatory attention cues induce systematic head rotations in the direction of upcoming lateral targets, thus implementing a behavioral mechanism to enhance sensory sampling of target sound.

Head rotations towards lateral targets were still evident when participants were instructed not to move, albeit reduced by approximately two orders of magnitude. This suggests that head rotations were planned but inhibited ^41^. This is reminiscent of microsaccades towards targets in covert visual spatial attention ^42–44^. Thus, head rotations associated with spatial attention are not fully suppressed on command, supporting the view of the premotor theory of attention ^15^ and the affordance competition hypothesis ^16^ that attentional orienting is tightly coupled to the preparation and competition of potential motor actions, even when overt movement is intentionally inhibited. This implies that planning and carrying out (miniature) head movements might relate to neural activity and/or task accuracy in typical, movement-constrained spatial attention investigations in the laboratory. However, this remains unexplained if head movements are not considered.

Permission to move the head improved target speech reception by ~3.3 %. The lack of an improvement for target speech on the right side was unexpected, but could be due to a right ear advantage for speech ^45,46^, which might mitigate the benefits of head rotation. Performance-beneficial head rotations to the left for targets and distractors on the left side suggest that listeners employed head rotations to enhance target sampling and segregate targets from distractors, respectively.

A hallmark of contemporary attention research is to separate target enhancement from distractor suppression ^3,47,48^. Convergingly, our findings suggest that head rotations are primarily employed to modulate sensory target processing. On the group-level, significant head rotations were found in the direction of lateral targets but not away from lateral distractors. The observed heterogeneity of head rotations in the case of lateral distractors might suggest that some listeners rotated their head towards lateral distractors to increase initial distractor sampling to support ensuing suppression ^49^, or to increase the signal-to-noise ratio on the ear closer to the target (i.e., better-ear listening) ^50,51^. Another group of listeners rather reduced the overall sampling of the distractor by turning the head away. Furthermore, when permitted to move the head, the number of errors associated with target-distractor confusions remained the same, indicating that head rotations do not help to suppress attention capture of the distractor ^52^. Instead, the permission to move improved accuracy by reducing reports of items that were neither targets nor distractors, suggesting that enhanced sensory sampling of the target reduced target uncertainty.

### Dissociable roles of neural alpha modulation and head rotation

The present study helps to understand the integration of goal-directed body movements with the neuroscientific basis of spatial attention. In line with movement-constrained laboratory studies, we found that spatial cue-induced lateralization of alpha oscillations reflects the focus of auditory spatial attention ^13,53,54^. Relatively reduced versus enhanced alpha power contralateral to a target versus distractor is thought to relate to neural enhancement versus suppression, respectively ^55^, but the link to body movements to targets or away from distractors is unclear. Analogous modulation of neural alpha oscillations in conditions that permit and exclude movement demonstrates that neural filtering does not depend on carrying out overt body movements.

Neural filtering and head rotation appear to contribute differently to spatial attention. Contrary to head rotation, alpha lateralization did not relate to improved task accuracy. Brain stimulation studies found evidence for causal contributions of lateralized alpha oscillations to auditory ^56,57^ and visual spatial attention performance ^58,59^. However, recent work casts doubt on the view that lateralized alpha oscillations modulate target versus distractor processing in early sensory areas ^60–62^, suggesting that spatial shifts of attention induce alpha modulation, not vice versa ^63^. Of note, significant alpha modulation in the present task occurred before the presentation of task-relevant and -irrelevant sounds, which might speak against a direct role in biasing neural representations of competing sounds. Rather, the present results appear to support a framework wherein lateralized alpha power codes the spatial positions of upcoming targets and distractors, which is followed by head rotations to enhance perceptual sampling of targets to improve task accuracy.

Recordings of neural activity alone reveal descriptions rather than explanations ^64^. Increased alpha power has been interpreted to reflect the intention to resist a contralateral distractor ^65^. Including recordings of head rotation, our findings suggest that this is only partly the case. A considerable number of participants aimed to separate the target from the distractor perceptually, thereby turning their heads towards instead of away from a lateral distractor. Thus, considering motor activity is inevitable if the goal is to explain the mechanistic basis of attentional filtering.

Furthermore, patterns of head rotation help understand the reference frame ^66^ of attention-induced modulation of parietal alpha oscillations. If alpha lateralization codes the spatial focus of attention with respect to momentary head direction (i.e., head-centered or egocentric) ^67^, head rotations should subsequently induce a modulation of neural filtering—which is not what we found. Instead, alpha modulation preceded head rotation and, importantly, retained its scalp-lateralized and temporal profile for conditions with and without permitted head movement. This result is rather in line with neural attentional filtering coding the focus of spatial attention in more allocentric or at least movement-invariant terms, with respect to the initial head direction at trial start.

### Coupled timing of neural filtering and head rotation

The primary objective of the present study was to examine the relationship between neural filtering and head rotation as a means of probing the mechanistic implementation of spatial attention. While no relation would suggest independent mechanisms, a negative relation would indicate *redundancy*, where stronger recruitment of head rotation reduces the need for neural filtering, and vice versa. Instead, we discovered a temporally ordered, positive association, suggesting a mechanistic chain from attention to action.

Modulation of lateralized alpha activity temporally preceded positively related changes of head rotation by 300–400 ms, corresponding to a delay of 3 to 4 alpha cycles. This speaks against the possibility that head movement-related activity drives lateralization of alpha oscillations. Given the dissociable roles of neural filtering and head rotations explained above, our results support an embodied model of attention ^6,68^ wherein a common upstream attention signal independently initiates neural alpha power modulation and head rotation, with the former reflecting the neural coding of relevant and irrelevant information in space and the latter modulating the sampling of sensory input to improve target processing.

Overlapping motor and attention systems in frontal cortex regions ^19,28^ might constitute a potential source of attention signals driving neural and motor activity. Furthermore, the superior colliculus is involved in shifting spatial attention ^69^ and contributes to head movements ^70^. Also, oculomotor activity might affect the interplay of neural activity and head rotation, as it relates to lateralized alpha oscillations ^30,31^ and precedes head rotation ^34,35^. However, in the present study, the HEOG related to temporally lagging head rotation but not to modulation of lateralized alpha activity. While these findings might suggest no central role of lateral eye movements in orchestrating the relation of alpha modulation and head rotation, it must be noted that the lack of eye-tracking data in the present study limits conclusions regarding the role of oculomotor activity. Especially, recent evidence suggests a strong association of microsaccades with M/EEG alpha activity modulation ^71–73^. Therefore, future investigations to study the basis of spatial attention in neural and motor signals should use comprehensive movement data recordings, including microsaccades, obtained through eye-tracking.

### Limitations of the study

The observed relation of head rotations to task performance might underestimate the impact of head rotations in real life to some extent. Our design implemented a limited number of spatial loudspeaker arrangements and thus induced stereotypical patterns of head rotation, which likely reduced the variance across (correct versus incorrect) trials. Furthermore, the present study focused on head rotations but arguably, other types of body movements that might relate to spatial attention [e.g., (micro-) saccadic eye movements, acceleration and movement of the upper body, etc.] have not been assessed. Future studies should thus include more comprehensive recordings of body movements.

## Conclusion

Spatial selective attention relies on the coordination of temporally coupled but functionally dissociable neural filtering and head rotation: Modulation of the parietal alpha rhythm codes spatial relevance, while head rotations implement overt sampling that improves performance by reducing target uncertainty. The temporally ordered coupling of neural alpha oscillations and over actions suggests coordinated control through a common upstream mechanism. This research invites models that treat overt movement as an integral component of cognitive control.

## STAR METHODS

Detailed methods are provided in the online version of this paper and include the following:

- KEY RESOURCES TABLE
- RESOURCE AVAILIBILITY
  - Lead contact
  - Materials availability
  - Data and code availability
- EXPERIMENTAL MODELS AND STUDY PARTICIPANT DETAILS
  - Participants
- METHOD DETAILS
  - Stimulus materials and task design
  - Individual stimulus adjustment
- QUANTIFICATION AND STATISTICAL ANALYSIS
  - Recording and analysis of head motion data
  - Recording and analysis of EEG data
  - Relative timing of neural filtering and head rotation
  - Statistical analyses

## Resource availability

### Lead contact

Further information and requests for data and code should be directed to the lead contact, Malte Wöstmann.

### Materials availability

This study did not generate new unique reagents.

### Data and code availability

- Data: Raw and preprocessed data have been deposited at OSF and are publicly available as of the date of publication. The DOI is listed in the key resources table.
- Code: All original code used for data analyses and figure generation has been deposited at OSF and is publicly available as of the date of publication. The DOI is listed in the key resources table.
- Additional information: Any additional information required to reanalyze the data in this paper is available from the lead contact upon request.

## Author contributions

Conceptualization: MW & JO; Data curation: MW; Formal analysis: MW; Investigation: MW; Methodology: MW & JO; Project administration: MW & JO; Writing-original draft: MW; Writing-review & editing: MW & JO.

## Declaration of generative AI and AI-assisted technologies in the writing process

During the preparation of the manuscript the authors used ChatGPT (GPT-5.2) to obtain more concise edits for certain single paragraphs at the draft editing stage. After using this tool, the authors reviewed and edited the content as needed and take full responsibility for the content of the published article.

## KEY RESOURCES TABLE

| REAGENT or RESOURCE | SOURCE | IDENTIFIER |
| --- | --- | --- |
| Deposited data |  |  |
| Raw and preprocessed behavioural and EEG data | This study | https://doi.org/10.17605/OSF.IO/BF4CU |
| Software and algorithms |  |  |
| Matlab R 2023b | Mathworks, Natick, MA, USA | https://de.mathworks.com/products/matlab.html<br>RRID:SCR_001622 |
| Psychtoolbox 3 | Brainard | http://psychtoolbox.org/<br>RRID:SCR_002881 |
| Palamedes Toolbox | Prins & Kingdom | https://www.palamedestoolbox.org/<br>RRID:SCR_006521 |
| Custom Matlab scripts | User-generated | https://doi.org/10.17605/OSF.IO/BF4CU |

### EXPERIMENTAL MODELS AND STUDY PARTICIPANT DETAILS

#### Participants

N = 33 human participants took part in the present study (26 females & 7 males; 32 right-handed & 1 left-handed; *M*_*age*_ = 23.15 years; *SD*_*age*_ = 2.97). Data of 7 additional participants were recorded but not used in the analysis due to missing data or technical problems. Participants provided written informed consent and were compensated for participation financially or with course credit. All experimental procedures were approved by the local ethics committee of the University of Lübeck.

### METHOD DETAILS

#### Stimulus materials and task design

Auditory stimuli were German numbers 1–9, spoken by a female voice ^74^. Numbers were shortened to 250 ms duration (using Praat 6.2.22) and overall sound intensity was equalized.

The cued auditory spatial attention task (Fig. 1) was adopted from previous studies ^13,75^. One loudspeaker (Genelec 8020D) was placed in front of the participant (0°; ~85 cm distance; above the monitor). The other loudspeaker was either placed on the left or right side (i.e., –90° or +90°; ~85 cm distance). On each trial, one loudspeaker was the cued target and the other was the distractor, implementing two conditions with lateral targets (*select-left, select-right*) and two conditions with lateral distractors (*suppress-left, suppress-right*).

Each trial started with a baseline wherein a fixation cross was presented (500–1000 ms, randomly jittered in 10-ms steps). Next, a visual spatial cue appeared (500 ms; arrow hat pointing left, right, or front). During the ensuing delay period, the cue changed to a fixation cross (1500– 2500 ms, randomly jittered in 1-ms steps). Then, the auditory stimulus was presented, which consisted of two different numbers, each presented at one loudspeaker location (front-left or front-right; duration: 250 ms). After stimulus presentation, the response screen appeared, showing a question mark. Participants had to report the number presented at the cued location using a number pad. There was no time out for the behavioral response.

Each participant performed 960 trials, separated in 8 blocks à 120 trials. In half of the blocks, participants were allowed to move their head, while movement was discouraged in the other half. The instruction to move / not move alternated from every block to the next. The loudspeaker setup (front–left vs. front–right) changed after every two blocks (participants with odd and even IDs started with front–left and front–right, respectively). For all trials in a block, the cue direction (front vs. side) varied randomly with the constraints that each cue appeared on half of the trials and that no cue would repeat more than three times. For every combination of setup (front–left vs. front– right) and movement (move vs. no-move) there were 2 blocks. For each participant, we randomly selected 240 pairs of target and distractor numbers and used these in randomized order for every combination of setup and movement.

The experiment was implemented using Psychtoolbox 3 ^76^ for Matlab (MathWorks, Inc., Natick, USA).

#### Individual stimulus adjustment

Before the main experiment, the individual signal-to-noise ratio (SNR) for target and distractor stimuli was determined to achieve accuracy of around 50 % (using the Palamedes toolbox for Matlab; Prins & Kingdom, 2018). Target numbers were presented in front and distractors either on the left or right side. A one-up, one-down adaptive staircase procedure was used ^78^. The staircase started with an SNR of +4 dB and was lowered vs. raised by 2 dB after each correct vs. incorrect response, respectively. The staircase terminated after 50 trials. The average SNR across the last four reversals was used for all conditions of the main experiment. Movement was discouraged during the stimulus adjustment phase.

### QUANTIFICATION AND STATISTICAL ANALYSIS

#### Recording and analysis of head motion data

Head motion and the electroencephalogram (EEG) were recorded with a mobile EEG system (Smarting PRO Indoor, mBrainTrain, Belgrade) and digitized at 500 Hz. Head rotation was measured in angular velocity (°/s) around three orthogonal rotational axes (yaw, pitch, roll). Additionally, acceleration along the three axes was assessed (m/s^2^), but not analyzed for the purpose of the present study.

To quantify horizontal head rotation to the left vs. right side, single-trial rotation around the yaw axis was extracted and averaged across trials, separately for experimental conditions. Other rotational axes (pitch & roll) were inspected as well but showed negligible variation as a function of experimental conditions. To calculate head orientation, angular velocity was integrated over time, using the function *cumtrapz* in Matlab. Of note, due to the placement of the amplifier box hosting the gyroscope on the neck, the gyroscope was somewhat tilted for most participants. Thus, absolute rotational angles around the yaw axis underestimate the true rotation to some extent.

We adopted a measure of towardness from previous research on gaze bias ^21^ to express single-subject head rotation toward versus away from lateral targets/distractors. Therefore, the contrast was calculated of angular rotation in trials with the lateral stimulus on the left versus right side (time window: 0.35–1.13 s) as a standardized *z*-value.

#### Recording and analysis of EEG data

The EEG was recorded at 32 electrodes (Ag/AgCl; 10-20 placement), using a common mode sense (CMS) reference and a driven right leg (DLR) ground. All electrode impedances were below ~10 kΩ. To ensure equivalent placement of the EEG cap, the vertex electrode (Cz) was placed at 50% of the distance between inion and nasion and between left and right ear lobes. For EEG preprocessing and analysis, we used the FieldTrip toolbox ^79^ and Matlab R2023b. Offline, the data were cut in epochs of –1 to +5 s around cue onset and filtered between 0.3 (high pass) and 100 Hz (low pass). An independent component analysis (ICA) was used to inspect and remove artefactual components based on component time courses, topographic maps, and spectra (average number of removed components: 17.21 of 32; SD: 3.88). It has been shown that a relatively small proportion of components reflects brain activity in mobile EEG datasets, which makes it necessary to discard a large number of components to allow functional interpretation of the data ^80^.

Next, 10 % of trials with largest ranges at any channel in the time interval –0.2 to +2.5 s around cue onset were removed and the data were re-referenced to the average of all channels. Trials were then split into experimental conditions and time frequency representations were derived using complex Fourier coefficients for a moving time window (fixed length of 0.5 s; Hanning taper; moving in steps of 0.04 s) for frequencies 1–80 Hz with a resolution of 1 Hz and 2-Hz spectral smoothing.

To quantify neural filtering, two previously established alpha modulation indices (AMI) were calculated on oscillatory power in the 8–12 Hz band ^13^. To quantify neural selection of lateral targets, we calculated AMI_selection_ = (Pow_select-left_ – Pow_select-right_) / (Pow_select-left_ + Pow_select-right_). To quantify neural suppression of lateral distractors, we calculated AMI_suppression_ = (Pow_suppress-left_ – Pow_suppress-right_) / (Pow_suppress-left_ + Pow_suppress-right_). AMIs were then contrasted for eight electrodes on the left (C3, T7, CP1, CP5, TP9, P3, P7, O1) vs. right hemisphere (C4, T8, CP2, CP6, TP10, P4, P8, O2).

#### Relative timing of neural filtering and head rotation

Changes in head orientation (i.e., head rotation in °/s) around the yaw axis were extracted for single trials. To compute single-trial alpha lateralization, preprocessed single-trial EEG data were high-pass– and low-pass–filtered at 8 and 12 Hz, followed by computation of the alpha band envelope (ENV; using the *envelope* function in Matlab) for eight electrodes on the left (LH) and eight electrodes on the right hemisphere (RH). Next, we averaged across electrodes within each hemisphere and calculated the AMI_single-trial_ = (ENV_LH_ – ENV_RH_) / (ENV_LH_ + ENV_RH_), followed by calculation of the first temporal derivative to quantify changes in alpha lateralization.

To detect lateral eye movements in the single-trial raw (unprocessed) EEG signal, we applied a 100-Hz low pass filter and baseline-corrected the data by subtraction of mean amplitude in the time interval –0.1 to 0 s. Then, we calculated the amplitude difference at electrode F7–F8 (see also Lins et al., 1993) to estimate the horizontal electrooculogram (HEOG), followed by computation of the first temporal derivative.

The three single-trial measures (head rotation, AMI_single_trial_, HEOG) were resampled to 100 Hz and extracted in the time interval 0–3 s, relative to cue onset. Pairwise cross-correlations between single-trial measures were calculated using the *xcorr* function in Matlab.

### Statistical analyses

Average proportion of correct responses per condition and participant were transformed to rationalized arcsine units (rau) and submitted to repeated-measures ANOVAs followed by post-hoc *t*-tests. For dependent-samples *t*-tests, we report Cohen’s *d* effect size; for repeated-measures ANOVAs, we report partial eta-squared (*η*_*p*_^*2*^). To test for significant neural filtering and head rotation across time, we applied cluster-based permutation tests ^82^ with 10,000 permutations on angular velocity and alpha lateralization indices (AMI), respectively. Cohen’s *d* effect sizes for cluster tests were derived from dependent-samples *t*-tests calculated on the data in the time intervals of significant clusters.

## Supplementary materials

**Figure S1.**
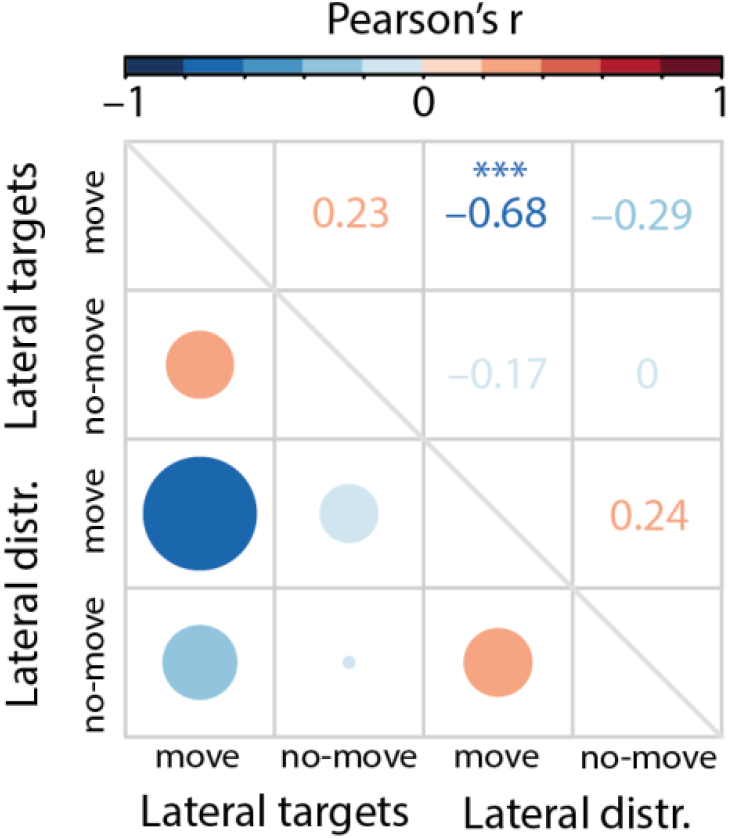
Heatmap shows Pearson’ r for pairwise correlations of single-subject head rotation (i.e., Towardness shown in Fig. 2D in the main manuscript). Note that only the correlation between the two movement conditions was significant (lateral targets, move & lateral distractors, move; *r* = –0.6794; *p* < 0.001), indicating that participants who moved more significantly in the direction of lateral targets also moved more significantly away from lateral distractors.

**Figure S2.**
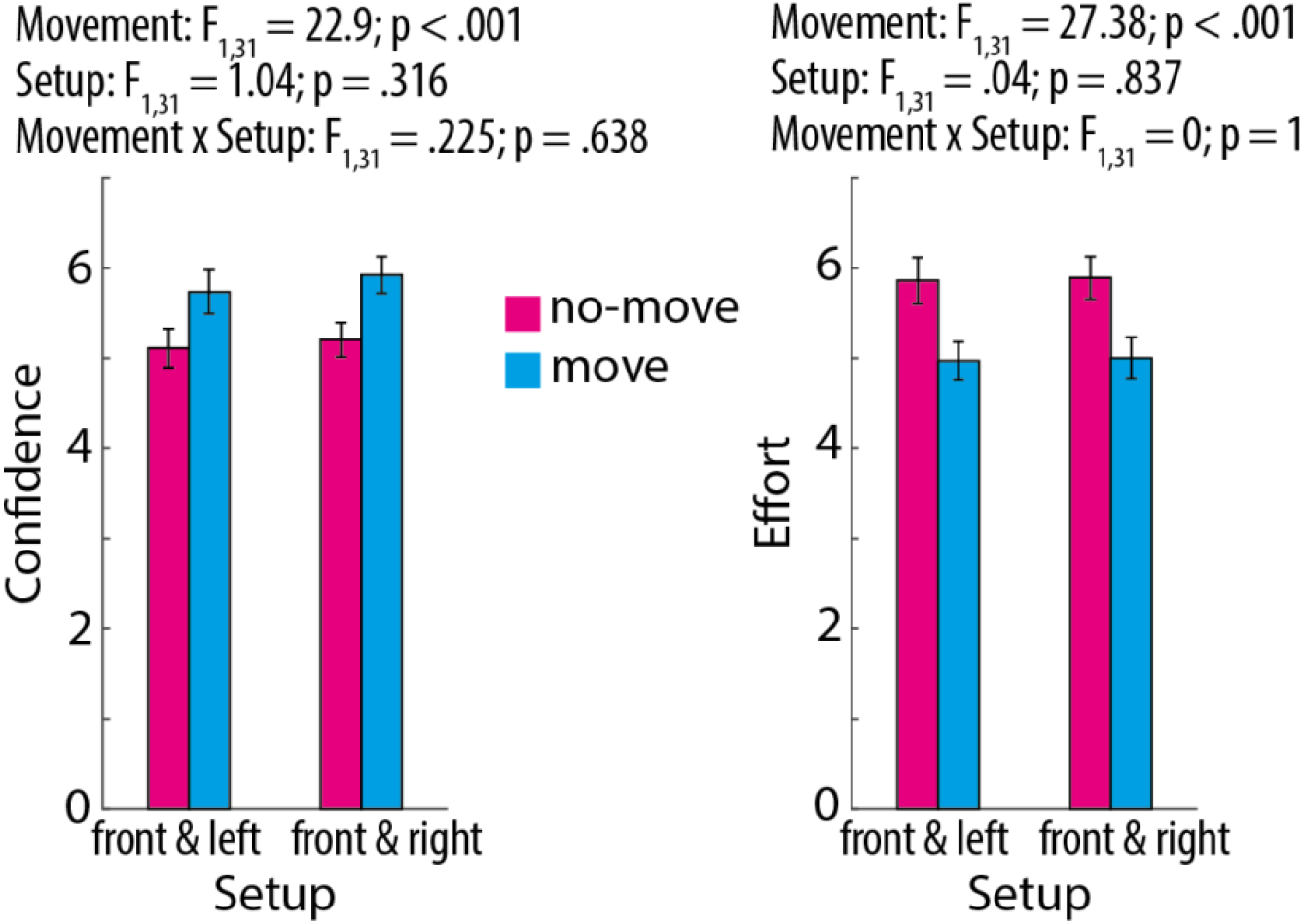
Confidence and effort ratings were collected in the end of each block on a scale from 1 (very low confidence / very low effort) to 10 (very high confidence / very high effort). Confidence was probed with the item “How confident are you that you reached a good performance level in this block?” Effort was probed with the item “How effortful did you experience the listening task in this block?” Results of two-way repeated measures ANOVAs with the factors movement (move vs. no-move) and loudspeaker setup (front & left vs. front & right) are shown above bar graphs. Bars show mean confidence/effort, error bars show ±1 SEM. Irrespective of the speaker setup, confidence ratings were significantly higher and effort ratings lower for blocks with movement permitted.

**Figure S3.**
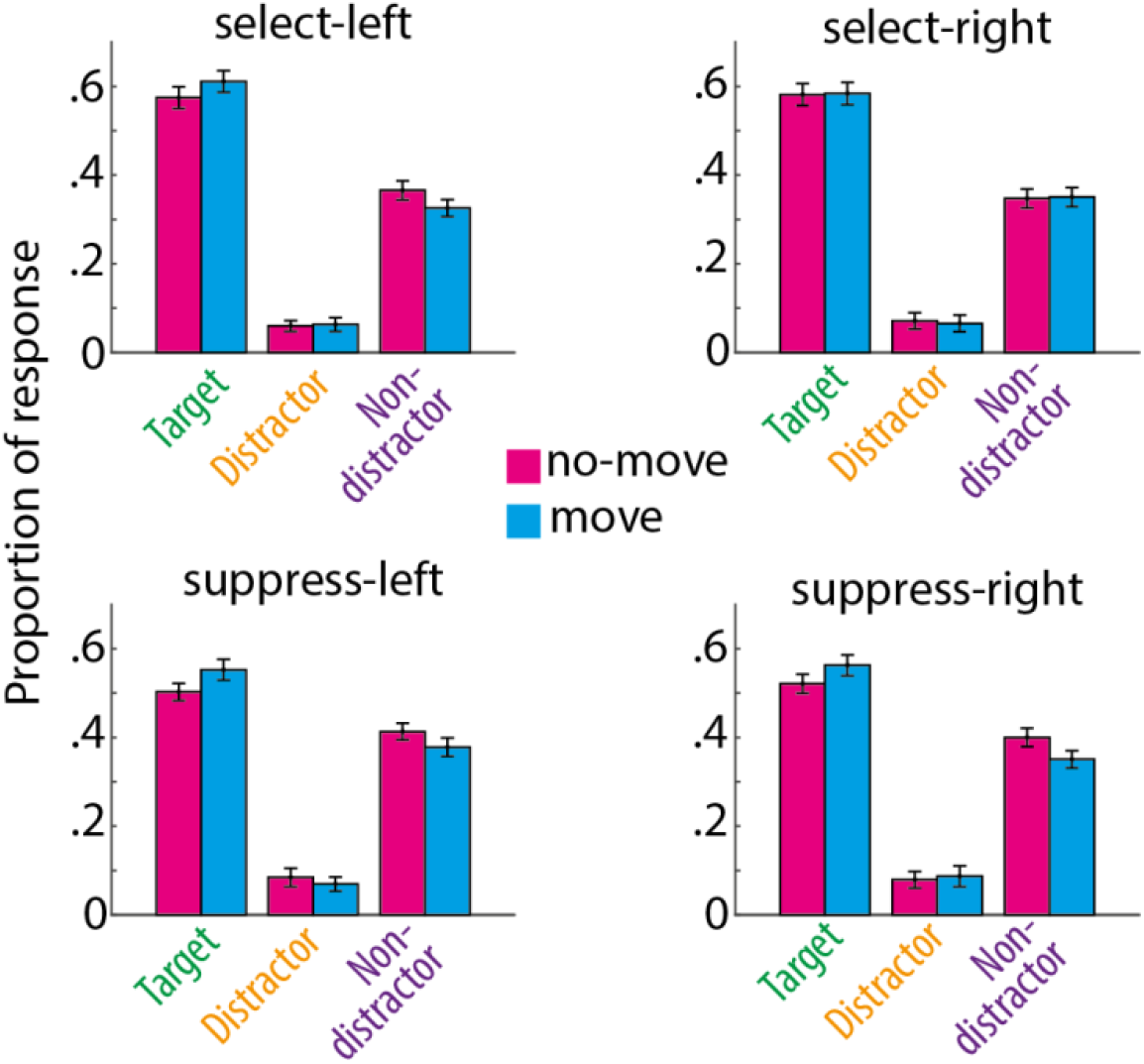
Bars show average proportion of responses for different response types for the four different experimental conditions. Error bars show ±1 SEM.

**Figure S4.**
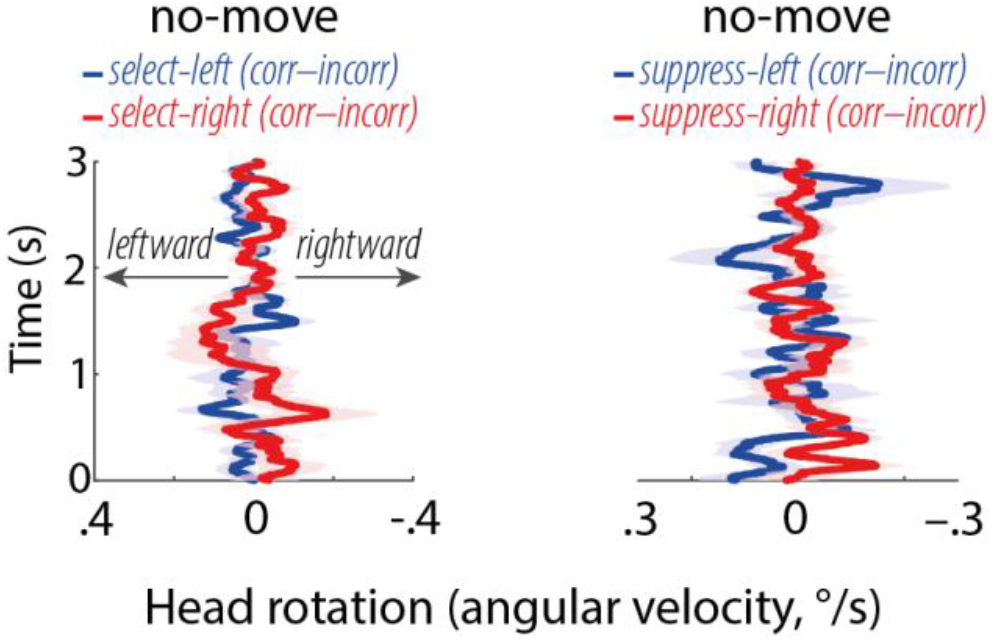
(**Left**) Lines and shaded areas show average angular velocity ±1 SEM for the difference of correct versus incorrect trials for trials with lateral targets and without movement permitted. (**Right**) Same as left but for trials with lateral distractors. No significant differences between correct and incorrect trials were found for these conditions (all *p* > .05, cluster-based permutation tests).

